# A hyperactive variant of the *Escherichia coli* anaerobic transcription factor FNR enhances ionizing radiation resistance

**DOI:** 10.1101/344382

**Authors:** Steven T. Bruckbauer, Joseph D. Trimarco, Elizabeth A. Wood, John R. Battista, Michael M. Cox

## Abstract

We have previously generated four replicate populations of ionizing radiation (IR)- resistant *Escherichia coli* though directed evolution. Sequencing of isolates from these populations revealed that mutations affecting DNA repair (through DNA double-strand break repair and replication restart), ROS amelioration, and cell wall metabolism were prominent. Three mutations involved in DNA repair explained the IR resistance phenotype in one population, and similar DNA repair mutations were prominent in two others. The remaining population, IR-3-20, had no mutations in the key DNA repair proteins, suggesting that it had taken a different evolutionary path to IR resistance. Here, we present evidence that a variant of the anaerobic metabolism transcription factor FNR isolated from population IR-3-20 can play a role in IR resistance. An FNR variant is unique to IR-3-20 and suggests a role for altered global metabolism through the FNR regulon as a means for experimentally-evolved IR resistance.

## Introduction

Bacterial species that do not display unusual levels of resistance to ionizing radiation can acquire such resistance by directed evolution [1-4]. However, a lack of advanced DNA sequencing technology prevented molecular characterization of evolved IR resistance in studies carried out in the previous fifty years. Over the past decade, we have generated IR resistance in the model bacterium *Escherichia coli* via directed evolution. Modern genomic sequencing methods have facilitated characterization of the evolved populations. We previously subjected four separate populations of *E. coli* to 20 cycles of ^60^Co irradiation (sufficient to kill up to 99.9% of the cells) followed in each cycle by survivor outgrowth. The 20 cycles resulted in large gains in IR resistance in all four populations, designated IR-1-20, IR-2-20, IR-3-20, and IR-4-20 [5, 6].

An isolate from population IR-2-20, CB2000, was previously characterized to identify the genetic alterations underlying the IR resistance phenotype [6]. The effort focused on mutations that were fixed in the population and affected genes or pathways altered in additional populations. Though seven mutations made at least a minor contribution, three mutations affecting DNA metabolism accounted for the majority of the IR resistance phenotype of CB2000. These were variants of (a) the DNA repair protein RecA (D276N), (b) the replicative helicase DnaB (P80H), and (c) the putative helicase YfjK (A152D) [6]. Reliance of CB2000 on three variant DNA metabolism proteins for IR resistance implicated a role for enhanced DNA repair in IR resistance. Indeed, biochemical characterization of the RecA D276N variant revealed novel activities consistent with repair of genomes fragmented by IR exposure [7].

Sequencing of isolates from each population revealed a striking trend: RecA, DnaB, and YfjK variants appeared in populations IR-1-20, IR-2-20, and IR-4-20 [5, 6]. This result suggested that, in addition to IR-2-20, both IR-1-20 and IR-4-20 rely at least in part on enhanced DNA repair for IR resistance. However, of seven isolates sequenced from population IR-3-20, none contained mutations in *recA*, *dnaB*, or *yfjK*. Protein variants common to IR-3-20 which appear in the same protein or pathway in at least one other population include those involved in DNA replication restart (PriA V554I), amelioration of reactive oxygen species (ROS) (GsiB L289P and RsxD V45A), cell wall metabolism (NanT F406S), and anaerobic metabolism (FNR F186I) [5, 6]. IR-3-20 also contains a unique intergenic SNP between the *clpP* and *clpX* genes (*clpP*/*clpX* int). Variants of the ClpP and ClpX proteins were also identified in population IR-1-20 [5, 6]. Of these pathways, only protein variants affecting ROS amelioration and cell wall metabolism have been previously associated with minor contributions to IR resistance [6]. Previous studies have suggested that extraordinary ROS amelioration play a major role in IR-resistance of the radioresistant bacterium *Deinococcus radiodurans* [8-11], suggesting that IR-3-20 may have evolved to ameliorate ROS produced by IR rather than enhance existing DNA repair pathways.

A variant of the anaerobic metabolism transcription factor, FNR, is unique to population IR-3-20, although a truncated FNR variant appears in an isolate from IR-1-20 further evolved for another 20 rounds of selection [6]. We now present evidence that altered global transcription through the FNR variant of IR-3-20, F186I, can make a substantial contribution to experimentally-evolved IR resistance.

## Results

### The FNR F186I variant enhances IR resistance

The F186I variant of FNR is the only variant of this protein detected in any of the four populations of *E. coli* exposed to 20 iterative cycles of IR selection via ^60^Co irradiation [5, 6]. Another FNR variant (an introduced stop codon at M157) appeared in a population derived from the evolved isolate CB1000, after twenty further cycles of selection [6]. We constructed a strain in which FNR F186I was placed in an otherwise wild type genetic background lacking the e14 prophage. As the e14 prophage was lost in all four populations very early in the directed evolution trial [5], a genetic background lacking e14 is the environment in which all other mutations detected in these populations operate. Thus, wild type strains lacking e14 have become the genetic background used as a standard comparator in any study involving these four populations. We refer to this background as Founder Δe14. At a dose of 3000 Gy, the FNR F186I variant increases IR resistance relative to Founder Δe14 by approximately 10-fold (Fig 1). This increase is comparable to that of the major IR resistance-enhancing single mutations from the IR-resistant isolate CB2000 [6].

**Fig 1.**
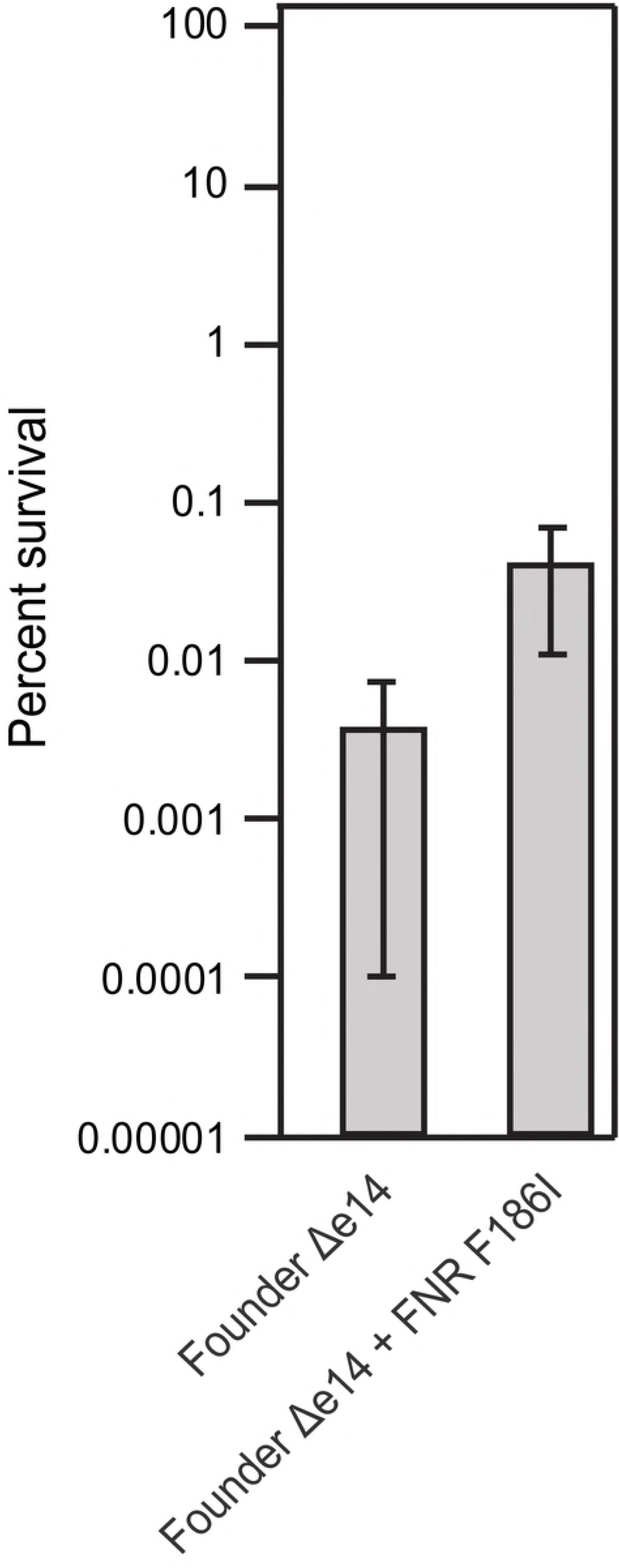
IR resistance of Founder Δe14 with the FNR F186I variant. The Y axis represents percent survival. Strains were assayed for survival of exposure to 3,000 Gy at exponential phase growth as described in the Materials and Methods section. The FNR F186I variant was moved from the IR-resistant isolate CB3000 into the Founder Δe14 background as previously described [6, 12]. The results represented indicate the average percent survival of 54 (Founder Δe14) and 23 (Founder Δe14 + FNR F186I) biological replicates. Error bars represent the standard deviation. The difference between the average of each strain is significantly different (p-value < 10^−12^) as calculated with a two-tailed student’s t-test. Raw data for Fig 1 is contained in Supporting Information (S1. IR resistance assays survival data).

### FNR F186I is a hyperactive FNR variant

We sought to better understand IR resistance derived from FNR F186I. FNR is a Fe-S binding transcription factor that regulates genes related to anaerobic metabolism [13, 14]. It has been previously reported that the F186 residue of FNR contacts the alpha C-terminal domain of RNA polymerase and that variants of residue F186 decrease activation of Class I and Class II FNR-dependent promoters [15, 16]. Therefore, we sought to determine the effects, if any, of the F186I variant on FNR-controlled transcription. We utilized a β-galactosidase fusion of the classical FNR-controlled *narG* promoter (P*_narG_*) to *lacZ* [17] to characterize the FNR F186I variant activity. We observed no difference in promoter activity in Founder Δe14 with the FNR F186I variant compared to Founder Δe 14 with the wild-type FNR F186 allele at early exponential (OD_600_ of 0.2) and stationary phase (overnight cultures grown for 15 to 18 hours) (Fig 2).

**Fig 2.**
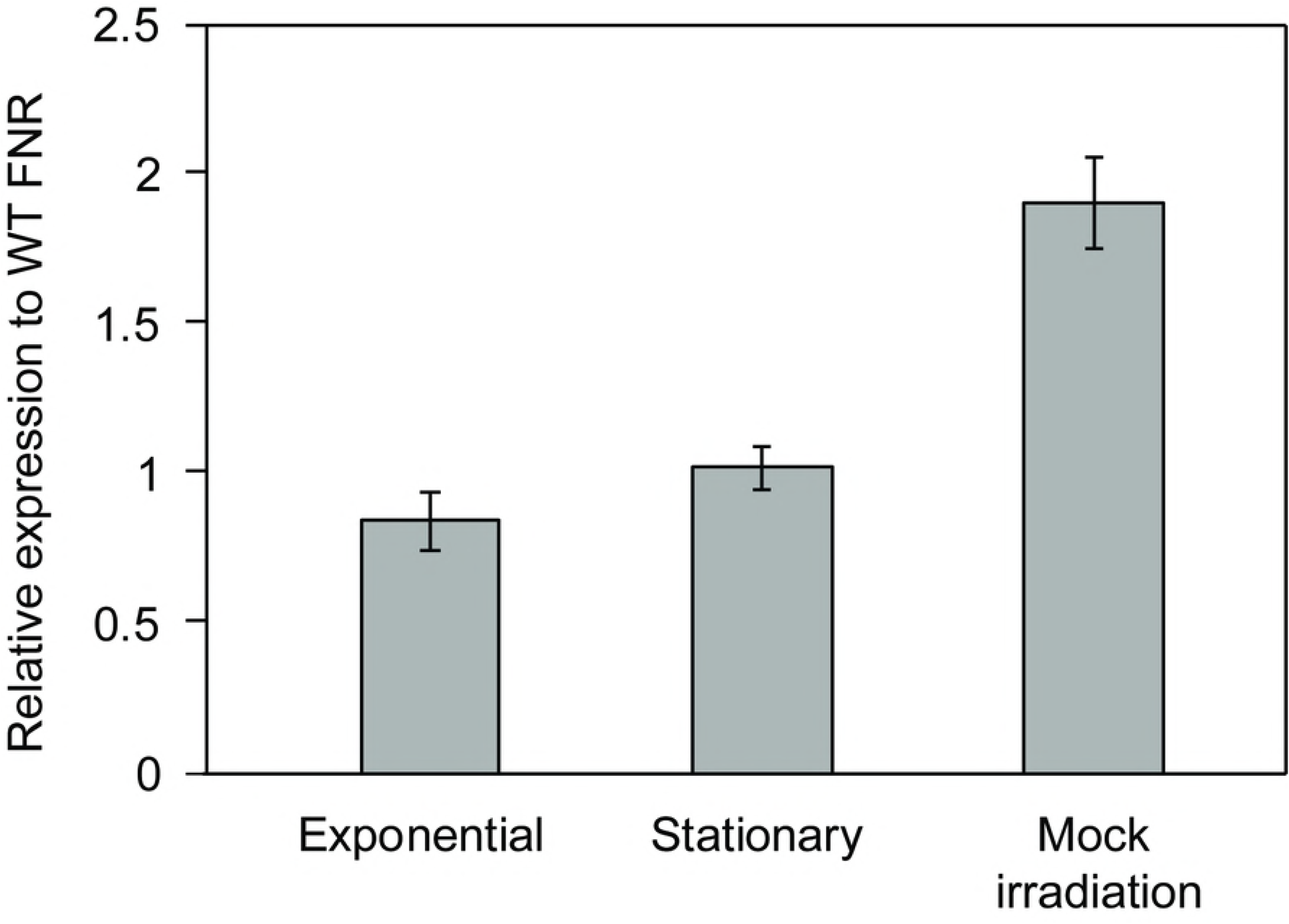
β-galactosidase activity of a P*_narG_*-*lacZ* fusion in Founder Δe14 + FNR F186I. β-galactosidase activity was assayed at exponential and stationary phase, and after a mock irradiation (exponential phase cells aliquoted in a 1.5 mL tube and incubated at room temperature for 8 hr). β -gal activity of Founder Δe14 + FNR F186I P*_narG_*-*lacZ* was normalized to the average β -gal activity of Founder Δe14 + P*_narG_*-*lacZ* (containing the wild-type FNR F186 allele) to determine relative expression. These data represent the results of two experiments of biological triplicates. The β-galactosidase assay was performed as described in the Materials in Methods section. The P_narG_-*lacZ* fusion has been previously described [17]. Data for these experiments are contained in the Supporting Information (S2. Beta-galactosidase raw data).

The conditions in which we irradiated strains (approximately an 8-hour duration for a dose of 3000 Gy, in sealed 1.5 mL tubes containing 1 mL of exponential phase culture) likely created a microaerophilic environment over time. A microaerophilic environment should increase the amount of active FNR dimers, which are destroyed by O_2_ [18, 19]. To determine the effect of these conditions on FNR activity, we performed a mock-irradiation, allowing culture to incubate at room temperature in 1.5 mL microfuge tubes as they do during irradiation assays. In mock-irradiation conditions, Founder Δe14 with the FNR F186I variant shows a 2-fold increase in transcription from the P*_narG_* – *lacZ* fusion compared to Founder Δe14 with the wild-type FNR (Fig 2). These results indicate that the FNR F186I variant allows for enhanced FNR activity under the conditions of our irradiation trials.

### FNR F186I enhances growth in rich medium without selection

The directed evolution protocol used to generate highly IR-resistant *E. coli* has an inadvertent, but useful, secondary selection step for enhanced growth during the outgrowth of irradiated survivors [5, 6]. To determine if the FNR F186I variant had any additional beneficial effect on growth, we utilized a previously described growth competition protocol without IR selection [6, 20]. After a duration of 48 hours, the growth competition assay revealed enhanced growth by Founder Δe14 with the FNR F186I compared to the parent strain (Fig 3). This is the first evidence that mutations selected for during our irradiation trials can both increase IR resistance as well as enhance growth without IR selection.

**Fig 3.**
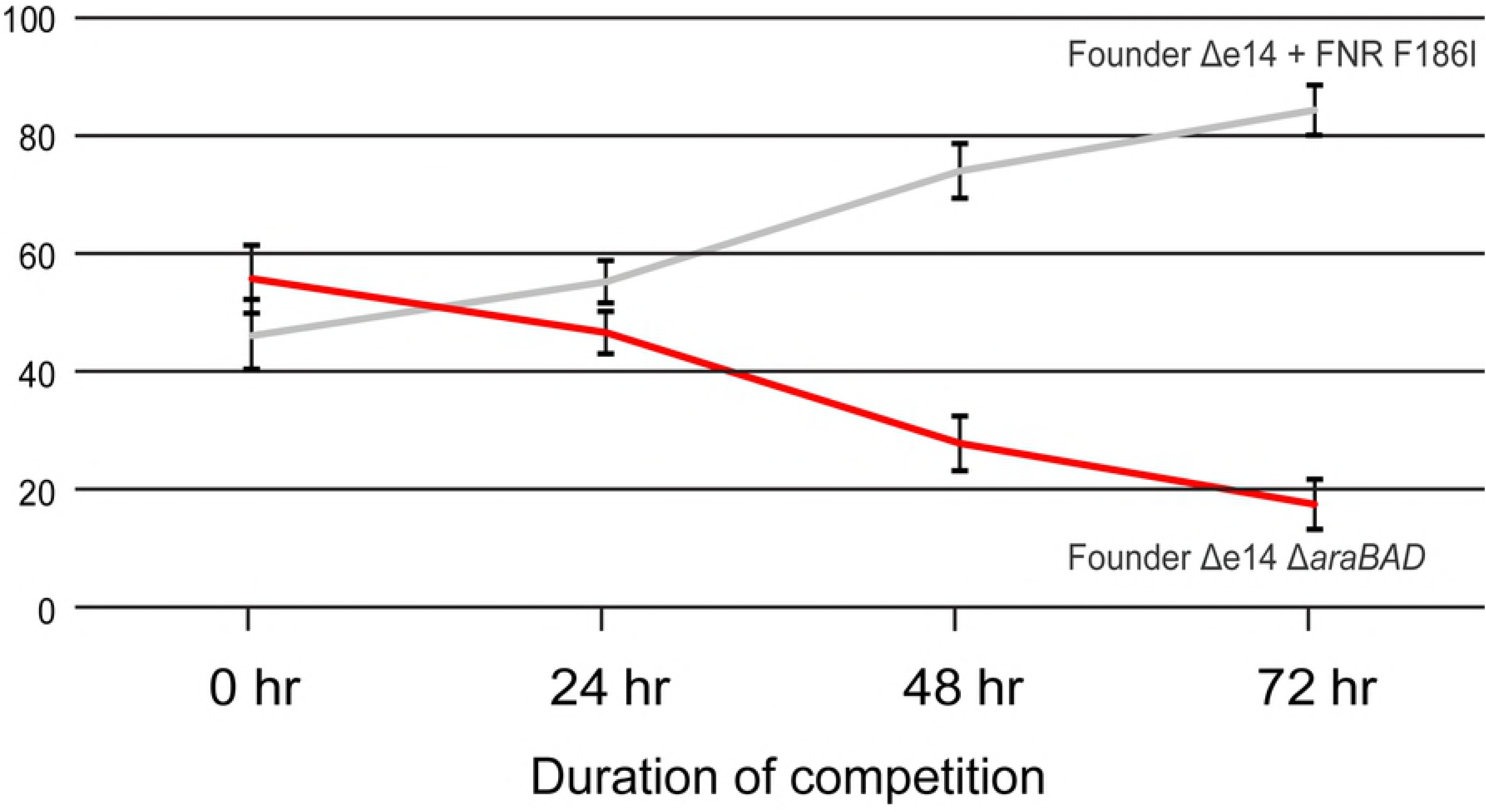
FNR F186I enhances growth competition of Founder Δe14 in mixed culture without selection. Founder Δe14 + FNR F186I outcompetes Founder Δe14. These data represent a single, representative competition against Founder Δe14 without the *araBAD* operon (a neutral mutation used to differentiate strains in the competition; loss of the *araBAD* operon results in red colonies on TA medium). Growth competitions were performed as described in the Materials and Methods. Data for these experiments are contained in the Supporting Information (S3. Growth competition raw data).

## Discussion

Utilizing four experimentally evolved populations of IR-resistant *E. coli* (IR-1-20, IR-2-20, IR-3-20, and IR-4-20) [5], we have begun to elucidate the means by which an naturally IR-sensitive organism can withstand extreme doses of IR. We have previously described experimentally-evolved IR resistance through enhanced DNA repair (the major contributors being RecA D276N, DnaB P80H, and YfjK A152D) [6]. Enhanced DNA repair appears to be a major mechanism of IR resistance in three of the four evolved populations, as IR-1-20, IR-2-20, and IR-4-20 each have prominent mutations affecting the RecA, DnaB, and YfjK proteins. IR-3-20 is unique in its lack of variants of these proteins, and the presence of an FNR variant. Here we demonstrate that this hyperactive FNR variant (F186I) affords IR-resistance, likely through a different mechanism than previously described DNA repair variants [5, 6]. Additionally, we present the first evidence that mutations which enhance IR resistance may also enhance growth without selection, indicating that selection cycles for IR-resistance also yielded selection for growth.

The downstream effects of the FNR F186I variant are unknown. Microarray analysis has implicated FNR as a regulator of 103 operons related to anaerobiosis, activating 68 and repressing 35 [21]. Of these FNR-controlled genes, *sodA* (which is repressed by FNR and encodes the Mn-binding superoxide dismutase) and *katG* (which is activated by FNR and encodes a bifunctional catalase/peroxidase) are two examples of genes that could (in principle) contribute to enhanced IR resistance through amelioration of IR-induced ROS.

The FNR F186I variant may also reduce endogenously produced ROS. Transfer of electrons from NADH and FADH_2_ to oxygen as a final electron acceptor in the electron transport chain is a major source of ROS [22]. FNR activates transcription of nitrate reductase A (*narGHJI* operon) and fumarate reductase (*frdABCD*), which allow for use of nitrate and fumarate as alternative electron acceptors. Enhanced production of nitrate reductase A and fumarate reductase due to the FNR F186I variant may thus allow cells to reduce levels of intracellular ROS, as nitrate and fumarate compete with O_2_ as a terminal electron acceptor. It is plausible that ROS-generation incidental to central metabolism may overwhelm already stressed ROS-amelioration mechanisms of irradiated cells, therefore reducing metabolic ROS may facilitate recovery from irradiation. Reducing endogenous ROS as means of IR resistance may parallel mechanisms used by *D. radiodurans*. It has previously been observed that in *D. radiodurans*, exposure to IR will alter gene expression to reduce metabolic ROS [23].

Why population IR-3-20 appears to have taken a different evolutionary path to IR resistance than the others is unclear; however, altered central metabolism through the FNR regulon appears to be a new potential contribution to experimentally evolved IR-resistance in *E. coli*. Further experimental evolution studies may elucidate the relationship between enhanced DNA repair and altered central metabolism, as these pathways may (or may not) appear within the same evolving populations over time.

## Materials and Methods

### Growth conditions and bacterial strains used in this study

Unless otherwise stated, *E. coli* cultures were grown in Luria-Bertani (LB) broth [24] at 37°C with aeration. *E. coli* were plated on 1.5% LB agar medium [24] and incubated at 37°C. Overnight cultures were grown in a volume of 3 mL for 16 to 18 hr. Exponential phase cultures were routinely diluted 1:100 in 10 mL of LB medium in a 50 mL Erlenmeyer flask and were grown at 37°C with shaking at 200 rpm and were harvested at an OD_600_ of 0.2, unless otherwise noted. After growth to an OD_600_ of 0.2, cultures were placed on ice for 10 min to stop growth before being used for assays.

Cultures were plated on tetrazolium agar (TA) for growth competition assays when noted [20].

All strains used for *in vivo* assays in this study are mutants of *E. coli* K-12 derivative MG1655 [25]. Strains are listed in the Supporting Information (S4. Strains used in this study). Genetic manipulations to transfer mutations or delete genes were performed as previously described [26, 27]. Lambda transductions to generate P*_narG_* – *lacZ* strains from parent strain RZ7350 [17] were carried out as follows. Incubated 10 mL of culture of the parent strain and the strains to be lysogenized overnight for approximately 18 hr at 37°C. To prepare the phage, added 300 µL to the turbid culture of the parent strain and shook at 200 rpm for 10 min at 37°C. Centrifuging for 10 min at 4700 x *g* pelleted the lysed cells, and the phage-containing supernatant was transferred to a new glass test tube. Five-hundred µL of the recipient strains were mixed with 6 mL of melted, 45 °C maximum, top agar and poured onto LB agar supplemented with 10 mM MgSO4 and 40 ug/mL X-gal. Once the top agar solidified, a 10 uL drop of prepared lambda phage was placed onto each plate. After the spot had dried, the plates were incubated overnight at 37°C. Blue plaques were streaked to isolation onto LB supplemented with 40 ug/mL X-gal until uniform, blue colonies were isolated. Lambda lysogens were confirmed as previously described [28].

### Serial dilutions and CFU/mL determination

All serial dilutions were performed in 1X phosphate-buffered saline (PBS) (for 1 L: 8 g NaCl, 0.2 g KCl, 1.44 g Na_2_HPO_4_, KH_2_PO_4_ 0.24 g with 800 mL dH_2_O, adjust pH with HCl to 7.4, then add remaining 200 mL dH_2_O). Unless otherwise stated, serial dilutions were performed with serial 1:10 dilutions of 100 µL of culture or previous dilution into 900 µL 1X PBS. Before transfer to the next dilution tube, samples were vortexed for 2 seconds and mixed by pipetting to ensure mixing. One-hundred µL of appropriate dilutions were aliquoted onto agar plates of the appropriate medium and were spread-plated utilizing an ethanol-sterilized, bent glass rod. For spot plating, 10 µL of each dilution was aliquoted onto agar plates of the appropriate medium and spots were allowed to dry before plates were incubated as in *Growth conditions*.

CFU/mL was calculated using the highest CFU count for each strain assayed that remained between 30 and 300 CFU (ex: 250 CFU on a 10^−4^ dilution plate would be used for calculation over 40 CFU on a 10^−5^ dilution plate).

### Ionizing radiation resistance assay

Cells from a single colony of each strain were cultured overnight and then grown to an OD_600_ of ~ 0.2 as in Growth conditions. One mL aliquots in 1.8 mL Eppendorf tubes were mixed by vortexing for 2 s and a 100 µL aliquot was removed and added to 900 µL PBS on ice as an initial 1:10 dilution for the non-irradiated control. Undiluted samples were then irradiated in a Mark I ^137^Cs irradiator (J. L. Shepherd and Associates) for a time corresponding to 3 kGy (~ 6.5 Gy/min). Irradiated samples as well as the non-irradiated samples were serial diluted and plated using typical methods. CFU/mL was determined for irradiated and un-irradiated samples, and CFU/mL was determined. Initial cell densities ranged from 1 - 6 x 10^7^ CFU/ml. For each experiment, each individual strain was tested in biological triplicate.

### β-galactosidase assay

Cultures to be assayed were prepared by incubating cells from a single colony of each strain overnight at 37 °C with aeration. This resulting overnight culture (grown 15-18 hr) was treated as the stationary phase culture to be assayed. Exponential phase cultures were grown by inoculating 10 mL of LB broth in a 50 mL Erlenmeyer flask with 70 µL of overnight culture and grown at 37 °C with shaking to an OD_600_ 0.2. These cultures were placed on ice for at least 5 minutes to stop growth before use. To perform a mock irradiation, two separate aliquots of 900 µL of exponential phase cultures in 1.5 mL Eppendorf tubes were inverted and incubated in the dark at room temperature (~24 °C) for 8 hr. Two aliquots for each replicate ensures that there is sufficient volume for 1 mL of culture to determine β-galactosidase activity and 100 µL to determine OD_600_.

The β-galactosidase assay was performed as follows. An OD_600_ reading was taken of each culture, and an appropriate amount (1 mL for exponential phase and mock irradiation cultures, and 50 µL for stationary phase) was aliquoted into 2 mL microcentrifuge tubes. Cells were pelleted via centrifugation at 6900 xg for 3 min, and supernatant was removed. Cells were resuspended in 1 mL Z buffer (0.06 M Na_2_HPO_4_, 0.04 NaH_2_PO4, 0.01 KCl, 0.001M MgSO_4_, to volume with purified dH2O), and 1 mL Z buffer was aliquoted in a 2 mL microcentrifuge tube for a blank sample. One-hundred µL chloroform and 50 uL 0.1% SDS were added to each tube. Each sample was then vortexed for 10 s and incubated at 4 °C for at least 10 min. Three samples at a time were removed from 4 °C and placed in a 28 °C water bath for 5 min. Two-hundred uL 4 mg/mL O-Nitrophenyl β-D-Galactopyranoside (ONPG) (Sigma-Aldrich, St. Louis, MO Cat#: N1127) dissolved in Z buffer was added to each sample. The development of yellow coloration for each sample was timed. Once the sample had become yellow (or after 30 minutes) the reaction was stopped by adding 500 uL 1M Na2CO3 and samples were placed on ice. All samples were spun for 10 min at 17000 xg in a microcentrifuge at 4 °C. One mL was removed from each sample, and the OD_420_ and OD_550_ was read. To determine β-galactosidase activity, the following equation was used where *t* is time of the reaction and *v* is the volume of culture used:

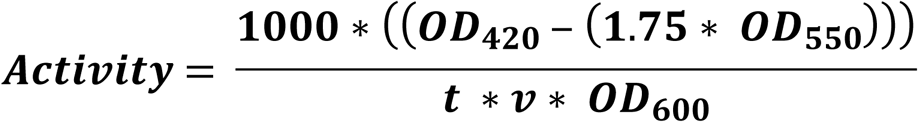

All β-galactosidase assays were performed using biological triplicate. To determine relative β-galactosidase activity compared to the parent strain, the activity of each mutant strain replicate was divided by the average activity of the wild-type triplicate in the given experiment.

### Growth competition assay

This assay was adapted from a previously published protocol [20]. To differentiate strains within the competition, a fitness-neutral deletion of the *araBAD* operon was introduced into one of the two strains. This deletion results in red colonies on tetrazolium arabinose (TA) agar plates (20). Briefly, overnight culture of each strain to be competed were mixed 1:1 in a 1.5 mL microcentrifuge tube. Samples were mixed by vortexing for 5 s and were serial diluted 1:10 in 900 µl phosphate-buffered saline (PBS) to a final dilution of 1:100,000. One hundred uL of the final dilution was spread plated onto TA agar plates to assay for CFU. Seventy µl of the remaining cell mixture was used to inoculate 5 mL of fresh LB media for growth overnight. This overnight culture was used to inoculate fresh media the following day, and 100 µl was serially diluted and plated as noted above. This procedure was repeated twice more over a period of two days. The number of white versus red CFU was noted after each day of the competition and the total percentage of the culture for each competitor was determined.

## Supporting Information

S1. IR resistance assays survival data

S2. Beta-galactosidase raw data

S3. Growth competition raw data

S4. Strains used in this study

